# Chimpanzees organize their social relationships like humans

**DOI:** 10.1101/2022.03.16.484664

**Authors:** Diego Escribano, Victoria Doldán-Martelli, Katherine A. Cronin, Daniel B.M. Haun, Edwin J.C. van Leeuwen, José A. Cuesta, Angel Sánchez

## Abstract

Human relationships are structured in a set of layers, ordered from higher (intimate relationships) to lower (acquaintances) emotional and cognitive intensity. This structure arises from the limits of our cognitive capacity and the different amounts of resources required by different relationships. However, it is unknown whether nonhuman primate species organize their affiliative relationships following the same pattern. We here show that the time chimpanzees devote to grooming other individuals is well described by the same model used for human relationships, supporting the existence of similar social signatures for both humans and chimpanzees. Furthermore, the relationship structure depends on group size as predicted by the model, the proportion of high intensity connections being larger for smaller groups.

## Introduction

Social network analysis has been a very active field for about a century, revealing the complex set of relationships that connect individuals (*1, 2*). Among the main objects of interest of social network analysis are personal or egonetworks, which consist of the social networks surrounding selected actors (*3*). A very general observation is that human egonetworks show a layered structure where each layer corresponds to relationships of different emotional closeness (*4– 6*). These layers have a definite emotional closeness: there is a layer of very close friends, a subsequent one of good friends, and so on. It is convenient to introduce the concept of nested circles, i.e., the sets of all the relationships up to a certain closeness. Typical circles established in the literature contain 5, 15, 50 and 150 individuals—with a scaling ∼ 3 between a circle and the next one. There is also evidence for a subsequent circle, formed by acquaintances, of about 500-600 people (*7*).

Social networks have also been studied in a diverse array of species, including mammals, birds, fish, amphibians, reptiles and invertebrates (*8–13*). In this context, the study of nonhuman primate social networks is of particular interest in light of the complexity of their societies, the variability between species, and their evolutionary proximity to us (*14*). Layered structures have been reported in both the distribution of primate social group sizes (*15*) and in groups of mammals living in multilevel social systems (mainly baboons, chimpanzees, elephants and dolphins) (*16,17*). However, while this suggests a possible connection to the layers of emotional closeness observed in humans, those results are not about individual egonetworks but about group-level social structures. To date, nobody has considered whether a continuous analysis of nonhuman primate social interactions reveals an underlying structure consistent with the structure that emerges in humans due to inherently limited resources of cognition and time.

To fill this gap, in this paper we present strong evidence that chimpanzees organize their relationships very much like humans do by means of a continuous version (*18*) of the theory introduced in (*19*), consisting of a resource allocation model based on two widely accepted assumptions: the capacity that an individual can invest in social relationships is finite, and relationships of a different intensity carry different costs. This mathematical approach allows us to advance our thinking beyond circles and assign a continuum value to a relationship, which is more reflective of real life and can include, for example, frequency of contact (*20*), number of messages exchanged (*21*), or duration of time spent together (*21*). The formalism developed in (*18*) was applied to face-to-face contact time (*22*), number of messages between Facebook users (*23*), and number of phone calls (*21*), showing a structure similar to that arising when intensities are regarded as discrete categories. This implies that it is not necessary to arbitrarily categorize the data to unveil its structure. As a consequence, egonetworks turn out to be characterized by a new universal scale parameter *η*, which plays the role of (and is consistent with) the scale factor ∼ 3 typically found in the discrete setup. We here apply the same formalism to grooming data extracted from over four years of observations of four groups of chimpanzees living in the Chimfunshi Wildlife Orphanage in Zambia, taking grooming as a proxy of the effort devoted to pairwise relationships. Grooming behavior is characterized by one individual manually or orally manipulating the hair or skin of another individual. While this behavior does serve a hygienic function, grooming is well-known to facilitate and reflect social bonding between individual chimpanzees (*24*). As we will see below, our results confirm that the time chimpanzees devote to grooming other individuals is well described by the continuous probability distribution predicted by the model, supporting the existence of similar social signatures for both humans and chimpanzees.

## Results

Generally speaking, the amount of grooming between primates is considered to be an indicator of their relationship quality (*25*). Therefore, we apply the formalism summarized above to grooming data of chimpanzees living in the Chimfunshi Wildlife Orphanage in Zambia between 2015 and 2019. At this sanctuary, chimpanzees live in four different populations without any interaction between individuals from different groups. The number of chimpanzees living in each group differs considerably (groups 1–4, *n* = 26, 60, 11, 14, respectively), and after preparing the data the number of individuals considered in each group for the analysis is reduced (groups 1–4, *n* = 21, 32, 10, 10, respectively). Full information about the chimpanzee population studied is provided in *Materials and methods*.

To analyze the data, we resort to the theoretical approach in (*18*), where the results for the discrete case (*19*) were extended to relationships characterized by continuous values. Briefly, the discrete approach assumes that *L* is the total number of relationships in an ego-network and *σ* is the average cognitive cost of a relationship. Relationships belong to *r* different categories, each of them bearing a different cost *s*_max_ = *s*_1_ > *s*_2_ > … > *s*_*r*_ = *s*_min_. As described in detail in (*19*) (see also *Materials and methods* below), using a maximum entropy approach it is possible to show that the number of relationships in one circle divided by the previous, smaller one, behaves approximately as

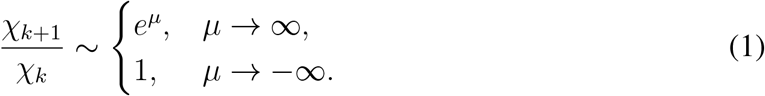

Where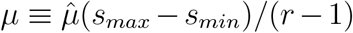, with 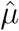 a Lagrange multiplier connected to the cognitive capacity constraint *σ*. Therefore, the circles satisfy an approximate scaling relation; in particular, for *µ* ≈ 1 the usual value of 3 found on empirical data is recovered. On the other hand, the theory also predicts a so-called ‘inverse’ regime, when *µ* < 0, in which most of the relationships are in the closest circle. This second behavior had not been described prior to the publication of (*19*), when it was checked against empirical data of small migrant communities, confirming its existence.

In the continuum case (*18*), circles are defined as the fraction of links *χ*(*t*) whose ‘distance’ to the individual is not larger than a specified value *t* (0 ≤ *t* ≤ 1). Their scaling ratio turns out to be controlled by a new parameter, *η*, the equivalent of *µ* (in fact, it can be shown that *η* ≈ (*r* − 1)(*e*^*µ*^ − 1)). In this continuous approach, the separation between the two regimes, the normal and the inverted ones, also takes place at *η* = 0. On the other hand, setting the number of circles to the usual value *r* = 4 and assuming, as empirically observed, that *e*^*µ*^ ≈ 3 (*5*), we then find *η* ≈ 6. More details can be found in *Materials and methods* below, and in the original references (*18, 19*).

With the above approach in mind, given a dataset of relationships with continuous weights, the scaling parameter *η* can be estimated using the maximum-likelihood method (see parameter estimation in *Materials and methods*). The type of fits of the function *χ*(*t*), giving the size of the continuous circles, to the data on individual chimpanzees is exemplified in Fig. 1 (plots of the fits for all the individuals considered in the study are provided in *Materials and methods*). The plots clearly show that the fits are not perfect, but on the other hand most points lie within the 95% confidence interval for the fitted distribution, and those that do not are not far from it. As can be seen in *Materials and methods* have been selected because they had more data for the fitting. One should bear in mind that the chimpanzee data are quite noisy because they have been obtained from chimpanzees living in large, naturalistic enclosures that lead to varying levels of animal visibility, and data have been collected by many different observers over the four-year period (see *Materials and methods*). Under those circumstances, the fits can be actually considered to be very good, and in fact if we compare them to those reported in (*18*), they are of a similar quality. Therefore, we can conclude that the continuum theory is a good description of how a chimpanzee distributes the time it devotes to grooming other individuals.

**Figure 1:**
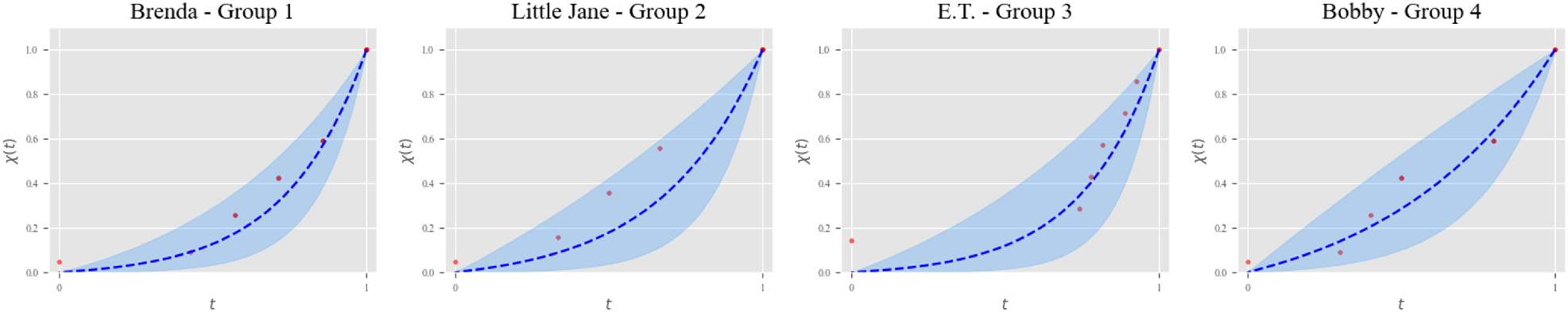
Examples of fittings for an individual of each group. Selected individuals are those for which there were more available data points. From left to right, individuals belong to groups 1, 2, 3, and 4. Shown in each case is *χ*(*t*), the fraction of links whose ‘distance’ to the individual is not larger than *t*. Red dots are actual data, representing the number of individuals who receive no more grooming than a fraction *t* of the maximum. The blue dashed line is the fitted function *χ*(*t*), and the blue-shadowed region is the interval of confidence.

Our analysis of the results for the parameters characterizing all the individuals studied is summarized in Fig. 2. The values for the parameter *η* obtained from the fits have a mode of approximately *η* = 4, while the mean value is close to the mode except for Group 2 where it is closer to 6. The range of obtained values for *η* falls within the range of expected values, and the mode indicates that typically the scaling ratio of the circles for chimpanzees would be somewhat smaller than for humans, except in Group 2. We believe that the reason for the difference of this group with respect to the others arises from the fact that it is distinctly larger than the rest, both pre- and post-filtering, and that this allows group members to develop a richer social life as far as grooming is concerned—i.e., it allows individuals to devote small intervals of time to grooming many others, leading to higher values of *η* and more low intensity relationships.

**Figure 2:**
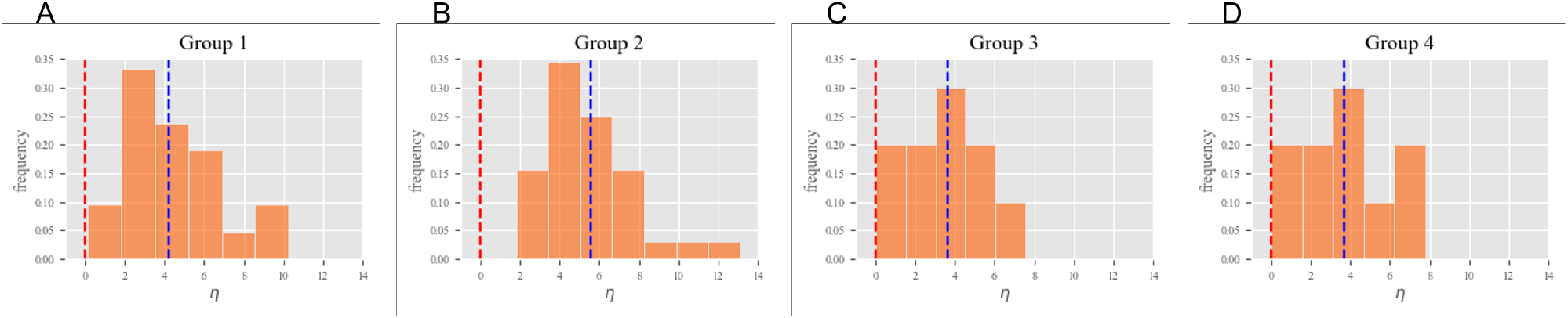
Histograms for *η* parameter distribution in each group. The red dashed line represents the change of regime *η* = 0 and the blue dashed line the mean value for each group.

The histograms presented show that there are no fits yielding negative values of *η*, which would indicate the existence of chimpanzees with an inverted structure of relationships, like those observed in humans (*19*). However, the values of *η* close to zero are on the border between regimes, so they should give rise to a larger fraction of individuals in the inner part of the distribution *χ*(*t*). Figure 3 confirms that this is indeed the case by showing the relationship structure and the *χ*(*t*) fitted function for two very different examples. On one side of the *η* scale we have the distribution of grooming times by Kit, whose *η* = 0.14 is rather typical of a structure that is intermediate between regimes. Kit devotes quite some time to grooming Kambo, Commander, Bobby and Val, and also a noticeable amount of time to other chimpanzees. Interestingly, this agrees with the fact that Kit is in group 4, the smallest one, in agreement with the situation in which the inverted regime is expected to arise: not so many possible individuals available to groom. On the opposite extreme of the *η* scale, Fig. 3 shows the results for Genny, in Group 1, with *η* = 10.3. In this case, we have a situation which is typical of the normal regime, with a lot of grooming devoted to her baby, some amount to her daughter Gonzaga, and very small amounts to many other individuals. Once again, this is to be expected in so far as Genny lives in Group 1, where there are many chimpanzees she can relate with. Therefore, the results corresponding to different values of *η* are also quite similar to those found for humans (*18*).

**Figure 3:**
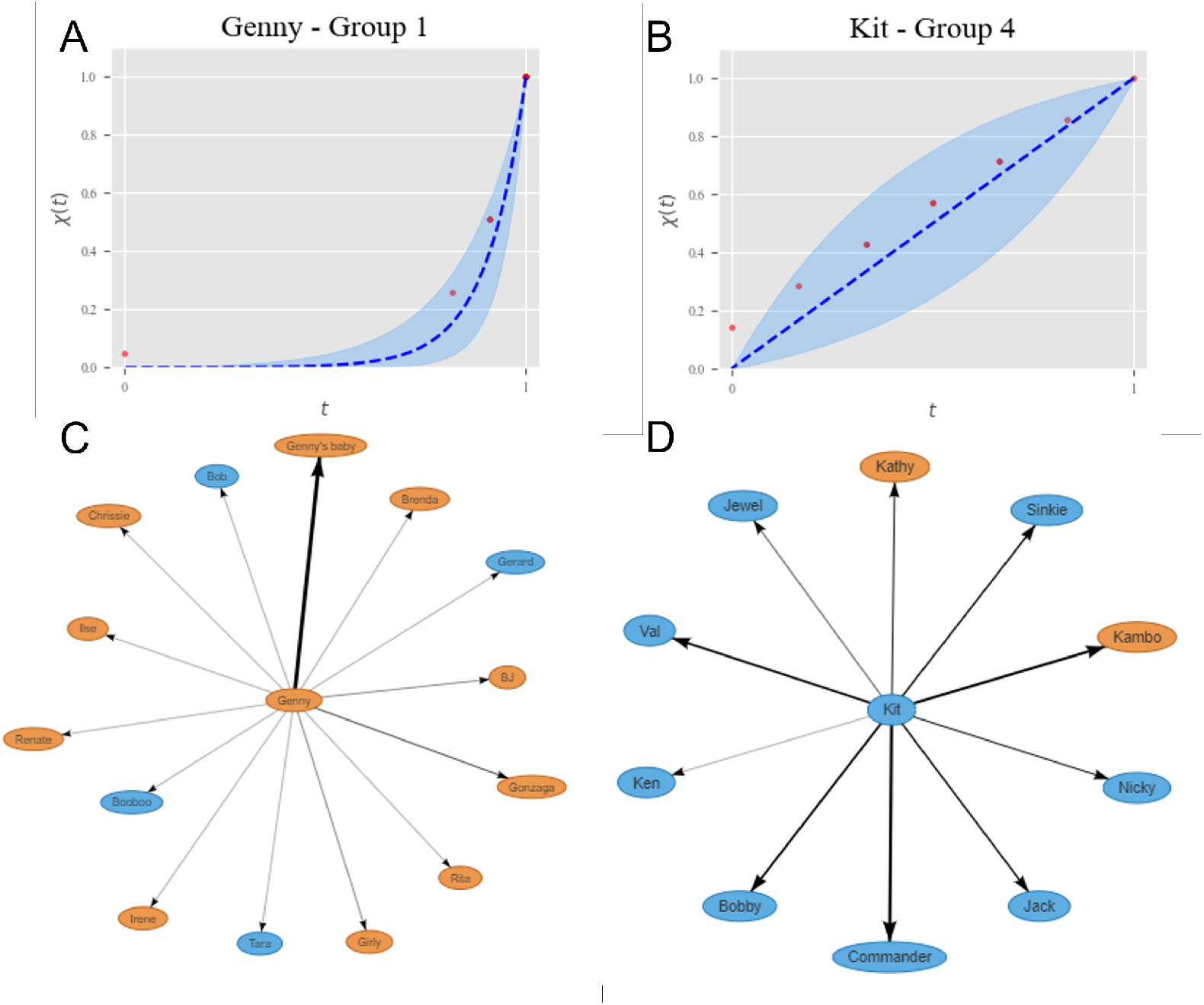
Examples of the relationship between the*η* parameter and the proportion of relationships of different intensity. *χ*(*t*) functions and egonetworks for Genny (Group 1 and *η* = 10.3) and Kit (Group 4 and *η* = 0.14). Arrows connect the focal individual with those it grooms, while the width of the arrows represents the total amount of time devoted to grooming a specific chimpanzee. Orange ovals represent females, blue ones represent males.

## Discussion

The results reported in this study show that chimpanzees appear to organize their grooming time in very good agreement with the prediction of the continuum resource allocation theory applied to humans in (*18*). In other words, chimpanzees distribute the time they devote to grooming other individuals in their group very much like humans organize their relationships when the relationship intensity is treated as a continuous value. These findings are in line with previous accounts that grooming can be considered a resource allocation problem (*26– 30*). As we see in humans, some chimpanzees solve this allocation problem by investing large amounts of grooming in a few other individuals (more so when the group is small), while others invest small amounts of grooming in many other individuals. In other words, they show the same two allocation strategies observed in human relationships and in the same situations, with inverted structures being more likely in small groups. We note that these findings corroborate grooming as an expression of friendship in non-human primates (*31*), yet more importantly indicate that different social strategies might be at play among chimpanzees, dependent on their immediate group structure. Our results show that chimpanzees living in larger groups employ their social capital differently than chimpanzees in smaller groups, like in humans (*19, 32*). This suggests that chimpanzees navigate their social environment flexibly, distributing their social resources across many group members when needed, yet investing more intensely in a few others when possible. In this light, any social forces that affect levels of group cohesiveness (e.g., to what extent a group forms a whole versus a modular, sub-grouping structure) indirectly shape a primate’s resource allocation strategy, a link we only know from the science of human sociology (*1*).

The novel finding here is that a nonhuman primate species, chimpanzees, appears to organize their affiliative relationships, or friendships, following the same pattern that we previously characterized for humans. Indeed, it appears that the mean group sizes of all primate species follow the same pattern as we see in the circles of human egonetworks (*15*). Furthermore, the global network structures of some primate species have a similar internal structure (*16, 17, 33*). By means of our continuum analysis of grooming data, we show that egonetworks in chimpanzees also exhibit a specific organization in terms of (the continuous equivalent of) circles. Our results thus establish that social networks in humans and chimpanzees show similar relationship structure. Further research could leverage the behavioral data of other primate species to determine whether the continuum analysis of egonetworks reveals a consistent pattern in other species living in large social groups.

## Materials and methods

### Environment description

At the Chimfunshi Wildlife Orphanage, chimpanzees live in large, forested enclosures ranging from 47 to 190 acres that consist of miombo grasslands and forests, a natural habitat for wild chimpanzee populations (*34*). In their enclosures, the chimpanzees have ample space to roam and exhibit species-typical behaviors, including engaging in natural fission-fusion dynamics (*35*). The four study populations at Chimfunshi live in separate enclosures which precludes the possibility for inter-group encounters. Apart from a small section between Groups 3 and 4, the chimpanzees from the different groups cannot see each other, yet live within hearing-distance from each other (i.e., the groups are stretched out over a crow-fly distance of 3km). Each of the four groups is composed of a mixture of wild-born chimpanzees and chimpanzees born at Chimfunshi. Wild-born chimpanzees come from various phylogenetic and geographic backgrounds, with a mixture of subspecies. The chimpanzees at Chimfunshi engage in natural foraging behavior on mostly fruiting trees, but also insects and small mammals present in their enclosures. Additionally, they are fed two times a day with a variety of fruits and vegetables to supplement their diets. At nights, the chimpanzees sleep in their woodland enclosures in self-constructed nests (for details see (*36*)).

### Information on the chimpanzees

**Table 1:**
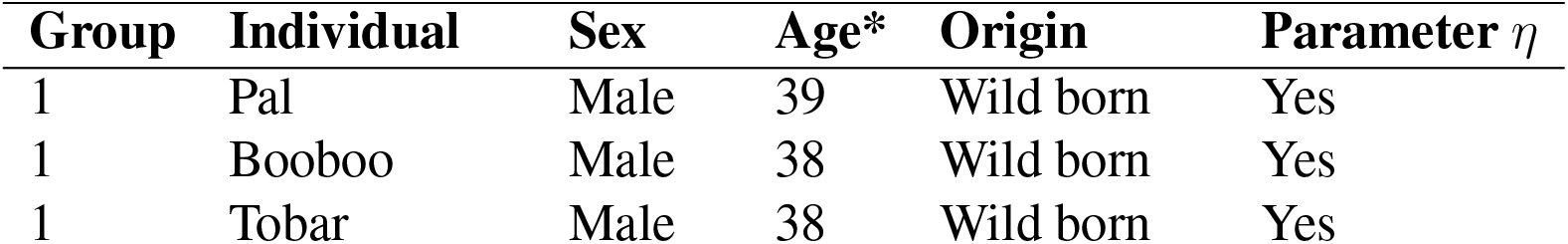
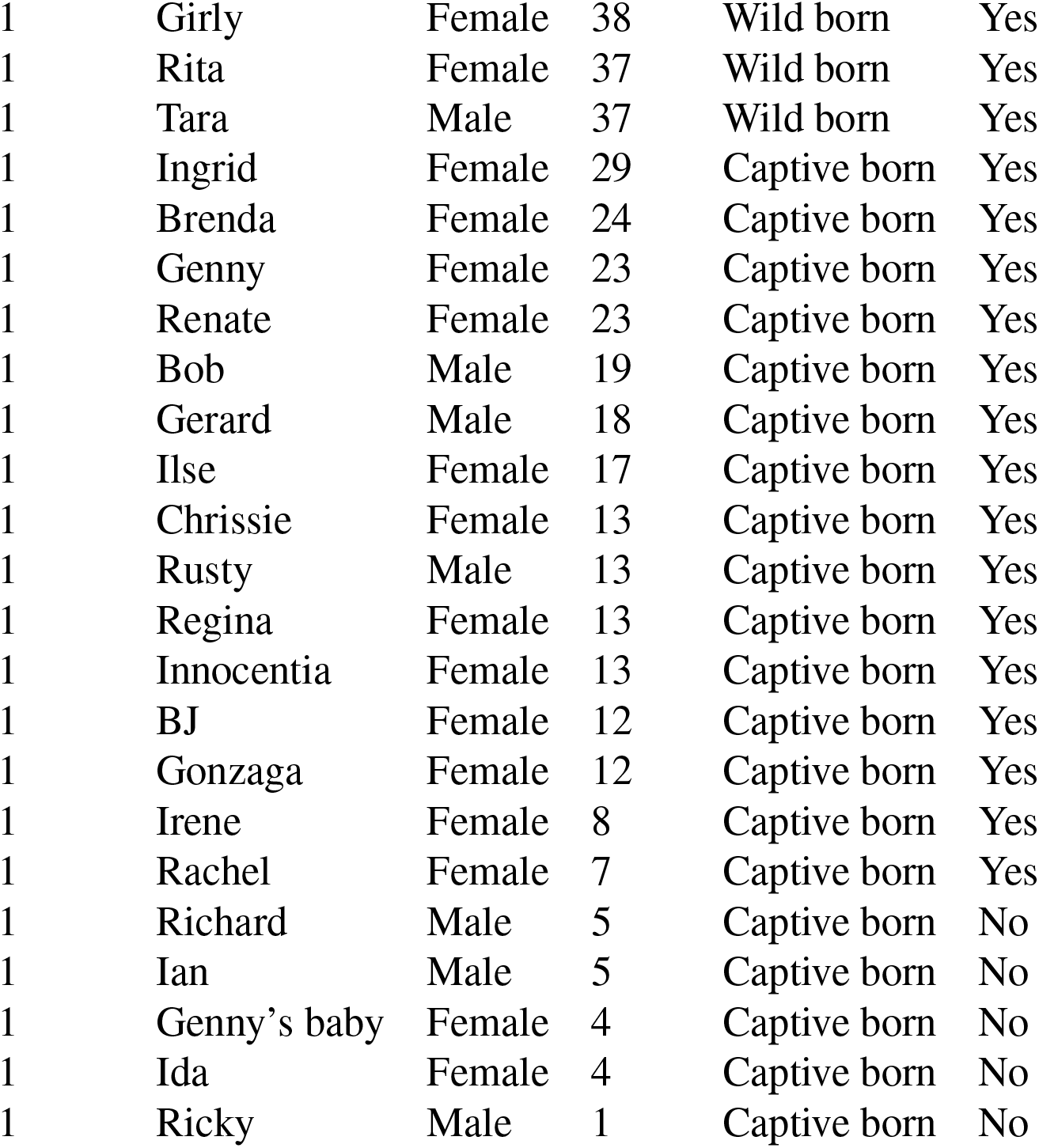
Group 1 info. *Age at the end of the observation window (late 2019)

**Table 2:**
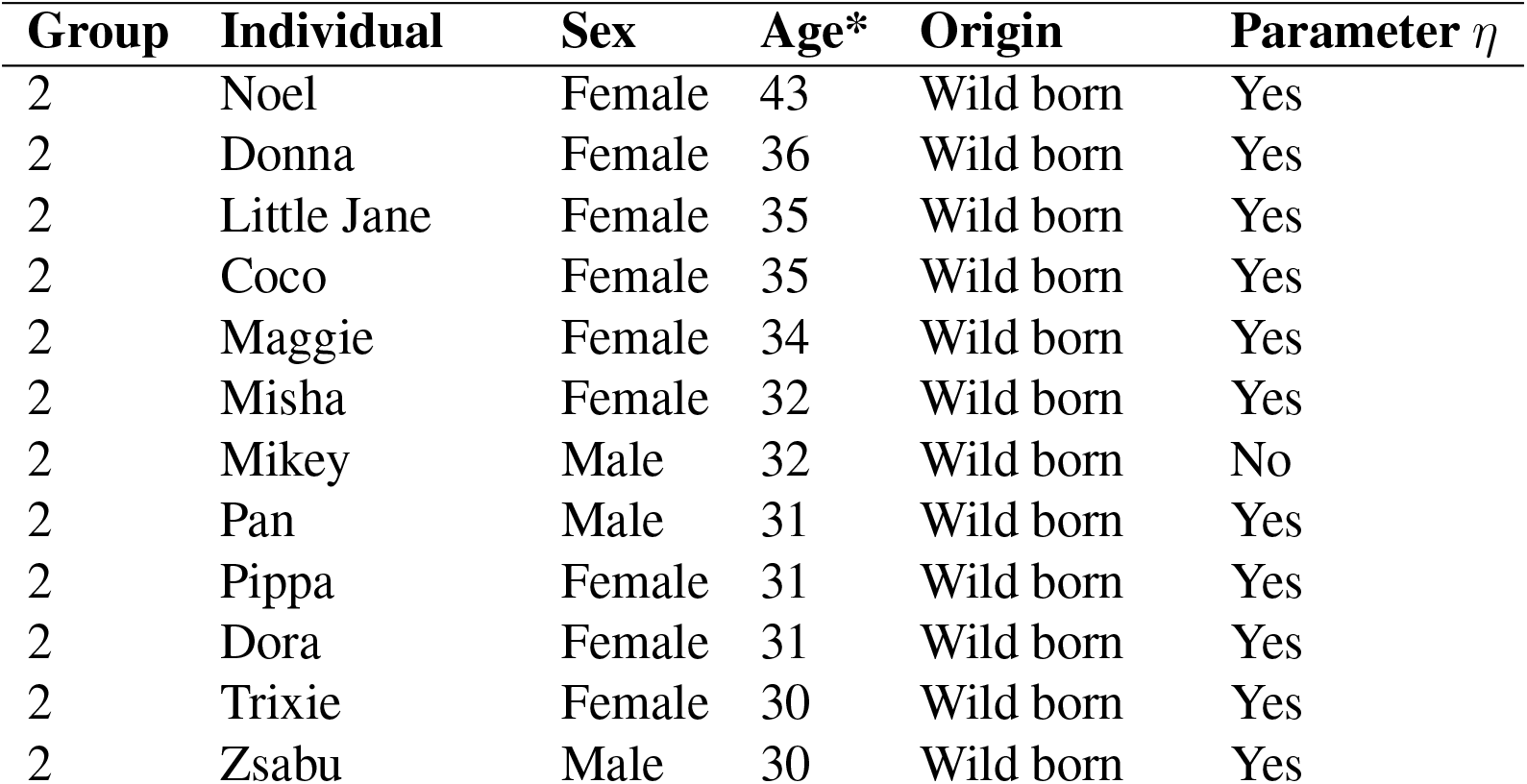
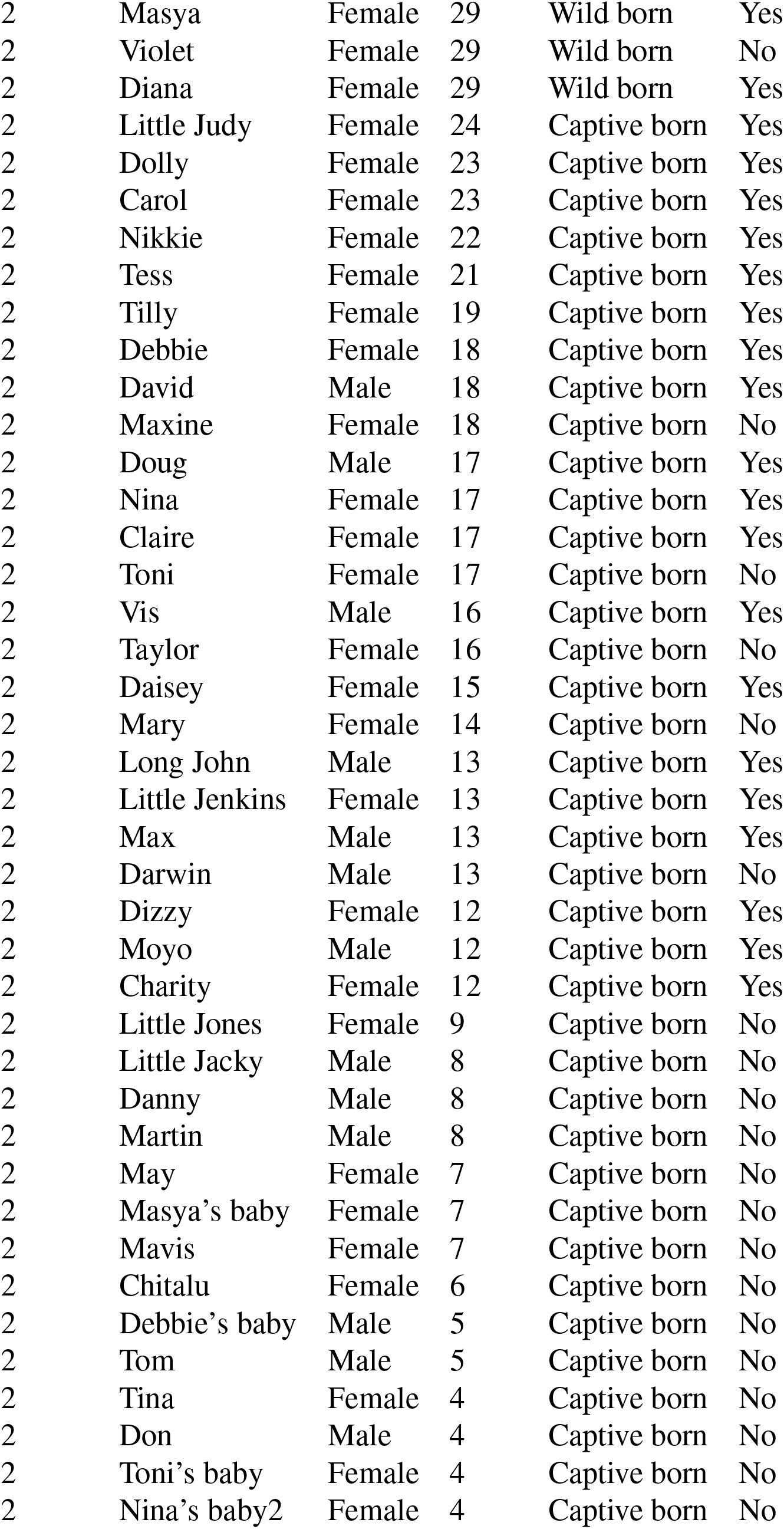
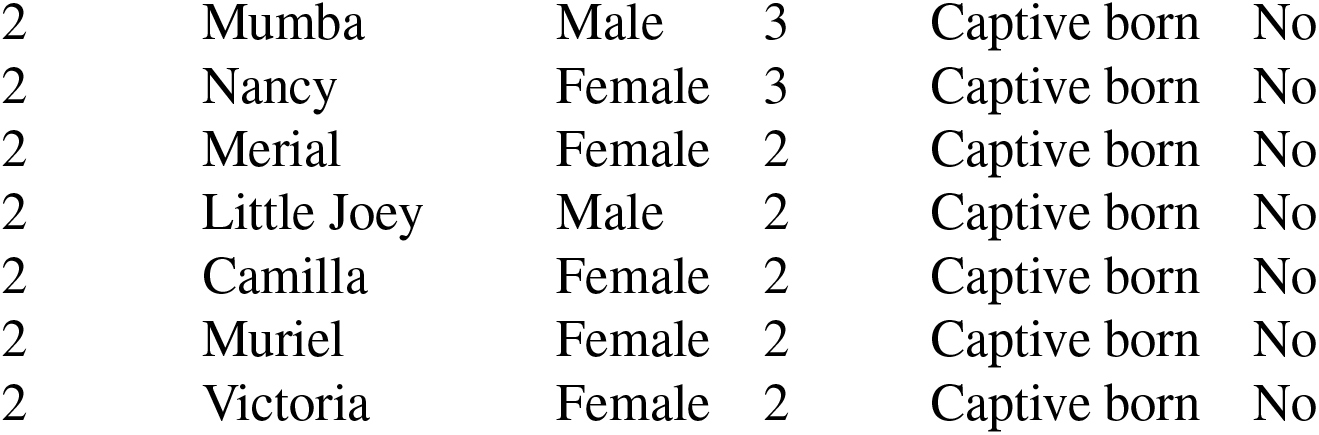
Group 2 info. *Age at the end of the observation window (late 2019)

| Group | Individual | Sex | Age* | Origin | Parameter $\eta$ |
| --- | --- | --- | --- | --- | --- |
| 2 | Noel | Female | 43 | Wild born | Yes |
| 2 | Donna | Female | 36 | Wild born | Yes |
| 2 | Little Jane | Female | 35 | Wild born | Yes |
| 2 | Coco | Female | 35 | Wild born | Yes |
| 2 | Maggie | Female | 34 | Wild born | Yes |
| 2 | Misha | Female | 32 | Wild born | Yes |
| 2 | Mikey | Male | 32 | Wild born | No |
| 2 | Pan | Male | 31 | Wild born | Yes |
| 2 | Pippa | Female | 31 | Wild born | Yes |
| 2 | Dora | Female | 31 | Wild born | Yes |
| 2 | Trixie | Female | 30 | Wild born | Yes |
| 2 | Zsabu | Male | 30 | Wild born | Yes |
| 2 | Masya | Female | 29 | Wild born | Yes |
| 2 | Violet | Female | 29 | Wild born | No |
| 2 | Diana | Female | 29 | Wild born | Yes |
| 2 | Little Judy | Female | 24 | Captive born | Yes |
| 2 | Dolly | Female | 23 | Captive born | Yes |
| 2 | Carol | Female | 23 | Captive born | Yes |
| 2 | Nikkie | Female | 22 | Captive born | Yes |
| 2 | Tess | Female | 21 | Captive born | Yes |
| 2 | Tilly | Female | 19 | Captive born | Yes |
| 2 | Debbie | Female | 18 | Captive born | Yes |
| 2 | David | Male | 18 | Captive born | Yes |
| 2 | Maxine | Female | 18 | Captive born | No |
| 2 | Doug | Male | 17 | Captive born | Yes |
| 2 | Nina | Female | 17 | Captive born | Yes |
| 2 | Claire | Female | 17 | Captive born | Yes |
| 2 | Toni | Female | 17 | Captive born | No |
| 2 | Vis | Male | 16 | Captive born | Yes |
| 2 | Taylor | Female | 16 | Captive born | No |
| 2 | Daisey | Female | 15 | Captive born | Yes |
| 2 | Mary | Female | 14 | Captive born | No |
| 2 | Long John | Male | 13 | Captive born | Yes |
| 2 | Little Jenkins | Female | 13 | Captive born | Yes |
| 2 | Max | Male | 13 | Captive born | Yes |
| 2 | Darwin | Male | 13 | Captive born | No |
| 2 | Dizzy | Female | 12 | Captive born | Yes |
| 2 | Moyo | Male | 12 | Captive born | Yes |
| 2 | Charity | Female | 12 | Captive born | Yes |
| 2 | Little Jones | Female | 9 | Captive born | No |
| 2 | Little Jacky | Male | 8 | Captive born | No |
| 2 | Danny | Male | 8 | Captive born | No |
| 2 | Martin | Male | 8 | Captive born | No |
| 2 | May | Female | 7 | Captive born | No |
| 2 | Masya's baby | Female | 7 | Captive born | No |
| 2 | Mavis | Female | 7 | Captive born | No |
| 2 | Chitalu | Female | 6 | Captive born | No |
| 2 | Debbie's baby | Male | 5 | Captive born | No |
| 2 | Tom | Male | 5 | Captive born | No |
| 2 | Tina | Female | 4 | Captive born | No |
| 2 | Don | Male | 4 | Captive born | No |
| 2 | Toni's baby | Female | 4 | Captive born | No |
| 2 | Nina's baby2 | Female | 4 | Captive born | No |
| 2 | Mumba | Male | 3 | Captive born | No |
| 2 | Nancy | Female | 3 | Captive born | No |
| 2 | Merial | Female | 2 | Captive born | No |
| 2 | Little Joey | Male | 2 | Captive born | No |
| 2 | Camilla | Female | 2 | Captive born | No |
| 2 | Muriel | Female | 2 | Captive born | No |
| 2 | Victoria | Female | 2 | Captive born | No |

**Table 3:**
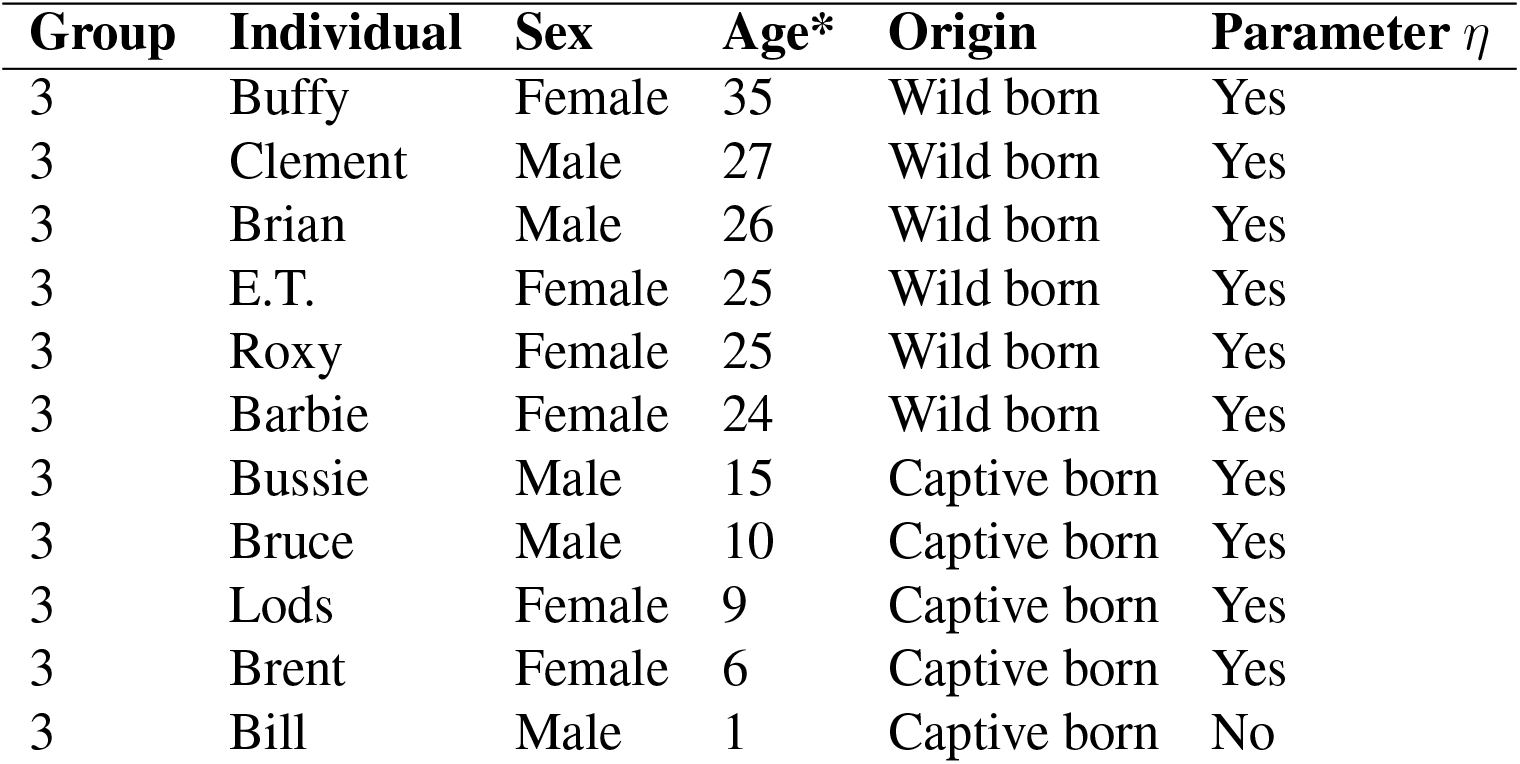
Group 3 info. *Age at the end of the observation window (late 2019)

| Group | Individual | Sex | Age* | Origin | Parameter $\eta$ |
| --- | --- | --- | --- | --- | --- |
| 3 | Buffy | Female | 35 | Wild born | Yes |
| 3 | Clement | Male | 27 | Wild born | Yes |
| 3 | Brian | Male | 26 | Wild born | Yes |
| 3 | E.T. | Female | 25 | Wild born | Yes |
| 3 | Roxy | Female | 25 | Wild born | Yes |
| 3 | Barbie | Female | 24 | Wild born | Yes |
| 3 | Bussie | Male | 15 | Captive born | Yes |
| 3 | Bruce | Male | 10 | Captive born | Yes |
| 3 | Lods | Female | 9 | Captive born | Yes |
| 3 | Brent | Female | 6 | Captive born | Yes |
| 3 | Bill | Male | 1 | Captive born | No |

**Table 4:** Group 4 info. *Age at the end of the observation window (late 2019)

| Group | Individual | Sex | Age* | Origin | Parameter $\eta$ |
| --- | --- | --- | --- | --- | --- |
| 4 | Nicky | Male | 29 | Wild born | Yes |
| 4 | Sinkie | Male | 26 | Wild born | Yes |
| 4 | Bobby | Male | 26 | Wild born | Yes |
| 4 | Kambo | Female | 24 | Wild born | Yes |
| 4 | Kathy | Female | 21 | Wild born | Yes |
| 4 | Miracle | Female | 20 | Captive born | Yes |
| 4 | Val | Male | 20 | Wild born | Yes |
| 4 | Commander | Male | 20 | Wild born | Yes |
| 4 | Kit | Male | 15 | Captive born | Yes |
| 4 | Jack | Male | 12 | Captive born | No |
| 4 | Leila | Female | 9 | Wild born | No |
| 4 | Ken | Male | 8 | Captive born | Yes |
| 4 | Jewel | Male | 6 | Captive born | No |
| 4 | Grace | Female | 5 | Wild born | No |

### Data collection

The grooming data were collected as part of a larger, ongoing data-collection effort at Chimfunshi aimed at assessing chimpanzee sociality over time (*36*). Trained staff members conduct focal follows daily with an every-minute scan sampling technique in the Zoo Monitor (ZM) application (*37*), a protocol which has been implemented and maintained by the authors (KAC, DBMH, EJCvL) since 2015. The protocol comprises 10min focal follows in which 10 scan points are scored. On each scan, all instances of proximity (¡1m), grooming, social play, and aggression by the focal individual are scored, including the identities of the interaction partners. The focal subjects are quasi-randomly selected as seen from the fence-line; the observation efforts start at a different location each day upon which the first-seen chimpanzee is chosen as focal. Subsequently, the next closest chimpanzee is chosen as focal. If the focal subject moves out-of-sight (which occurs because the enclosures are sizable), observers note the animal as out-of-view. Observation sessions last one hour on average.

### Data curation

We chose to focus on grooming behavior because of the established relationship between grooming behavior and dyadic relationship quality (*24*), and because we judged the grooming data to be most reliably and consistently collected over the four year period. For our analysis we consider a grooming interaction when an individual has been observed grooming another in the 10 minutes interval it acts as a focal subject, regardless of whether this action takes place during the whole of this period or only during a fraction of it. In this way, we can unify the criteria for considering a grooming action and reduce uncertainty in the data, as in many cases it starts or ends outside the focal observation period. As an alternative, we have assigned a weight to each grooming bout given by the number of minutes it lasted within the observation window. Data analysis using this criterion yields results in very good agreement with those presented here, so we retained the first procedure as is standard in the field.

Prior to obstructing networks, we filtered chimpanzees that have groomed less than five individuals in the study period, although we still consider the grooming actions performed on them. This condition does not influence the conclusions, since the calculus of the parameter *η* is performed individually for each chimpanzee, and is based on the fact that five is the size of the core of grooming egonetworks in primates (*15*). With this restriction, we basically select for the analysis of the egonetwork the chimpanzees older than nine years old, while the infants and the chimpanzees who died between 2015 and 2019 are not considered. Therefore, the restriction allows us to homogenize the population of chimpanzees studied and to extrapolate the results obtained to the case of adults.

### Theoretical background

For the sake of clarity in what follows, let us briefly summarize the main results from theoretical approaches to the circle structure. In the discrete case (*19*), it is assumed that *L* is the total number of relationships in an ego-network and *σ* is the average cognitive cost of a relationship. Relationships belong to *r* different categories, each of them bearing a different cost *s*_max_ = *s*_1_ > *s*_2_ > … > *s*_*r*_ = *s*_min_. As described in detail in (*19*), using a maximum entropy approach it is possible to obtain the probability that a given relationship of the ego-network belongs to category *k* as

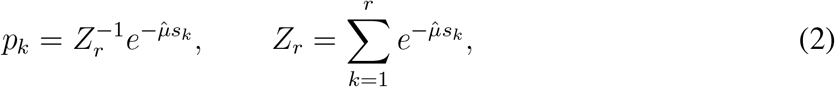

where 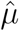 is fixed by letting *σ* be the expected cost *σ* = 𝔼 (*s*_*k*_). Using this probability distribution we can calculate *χ*_*k*_, the expected number of relationships with costs larger than or equal to that of category *k* (i.e., the size of the social circles, with *k* = 1 corresponding to the innermost one), as

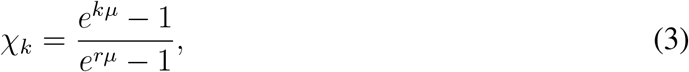

Where 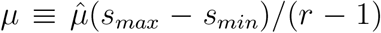 . As mentioned in the main text, it can subsequently be shown that, for large values of *µ*, the scaling ratio, i.e., the size of one circle divided by the previous one, behaves approximately as

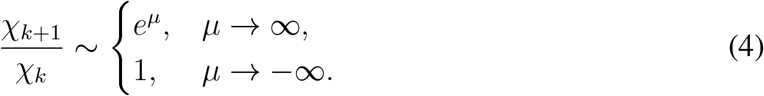

As discussed in (*19*), this result predicts the known regime for values of *µ* > 0, in which the circles satisfy an approximate scaling relation; in particular, for *µ* ≈1 the usual value of 3 found on empirical data is recovered. On the other hand, it also predicts a so-called ‘inverse’ regime, when *µ*< 0, in which most of the relationships are in the closest circle. This second behavior had not been described prior to the publication of (*19*), when it was checked against empirical data of small migrant communities, confirming its existence.

In the continuum approach (*18*) the key parameter is called *η*, and it is related to the average cost *σ* by the implicit equation

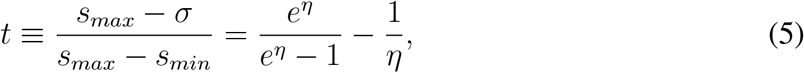

and thus *η* is actually a function *η*(*t*), with *t* defined in the equation above representing a normalized measure of the cost of a relationship (*t* = 0 corresponding to the highest cost and *t* = 1 to the lowest one). Once *η* is determined, the fraction of relationships with a normalized cost not larger than *t* is given by

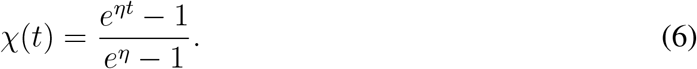

This is the curve that should fit the data. Notice that each individual will be characterized by its own value of *η*.

The scaling ratio of the circles can be obtained from the asymptotic behavior, for large *η*, of the logarithmic derivative of *χ*(*t*), the fraction of links whose ‘distance’ to the individual is not larger than *t*, which turns out to be

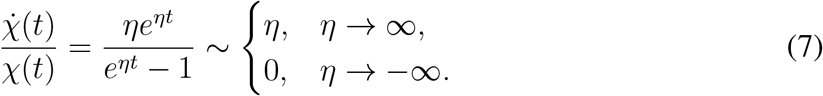

In this approach, the separation between the two regimes, the normal and the inverted ones, also takes places at *η* = 0.

Finally, to connect the two formalisms, we can use the fact that the discrete version of the left-hand side is (*χ*_*k*+1_ −*χ*_*k*_)/*χ*_*k*_Δ*t*; then, a comparison between (4) and (7) in the ordinary regime leads to *η*Δ*t*≈ *e*^*µ*^ − 1. Since Δ*t* ≈ (*r* − 1)^−1^, we obtain the equivalence

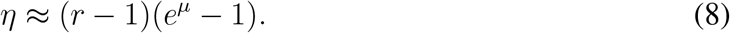

Interestingly, this result shows that the value of *µ* in the discrete model depends on the total number of layers, *r*. This fact had not been noticed in previous research because of the implicit assumption of the existence of *r* = 4 layers in the structure of egonetworks *(5)*. Setting *r* = 4 in (8) and assuming, as empirically observed, that *e*^*µ*^ ≈ 3 (*5*), we then find *η* ≈ 6.

### Parameter estimation

With the above approach in mind, given a dataset of relationships with continuous weights, the scaling parameter *η* can be estimated using the maximum-likelihood method. As described in (*18*), such an analysis leads to an expression equivalent to (5) to connect the range of data weights to the theoretical parameters, *η* and *σ*. Thus, for an empirical dataset we can find the values of *s*_*max*_ and *s*_*min*_, which are the largest/smallest possible costs an individual can invest in a relationship, respectively. Then, the value of *σ*, the total cost per item, is determined by

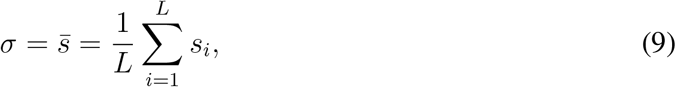

where *s*_*i*_ are the costs associated to each of the relationships, measured in the same units as *s*_*max*_ and *s*_*min*_, and *L* is the total number of relationships that an individual has. Once these variables are set, the parameter *η*, that characterizes the structure of the egonetwork of each individual, can be estimated solving (5) numerically. Furthermore, an expression for the 1 − 2*δ* confidence intervals associated to the parameter *η* can be found (see (*18*) for details). In what follows we choose a 95% confidence interval using *δ* = 0.025.

### Code implementation and availability

All the numerical analysis carried out are an adaptation of those used in (*18*) (available at https://github.com/1gnaci0/continuous-resource-allocation), using the Python packages scipy.optimize and scipy.integrate.

### Grooming networks

**Figure 4:**
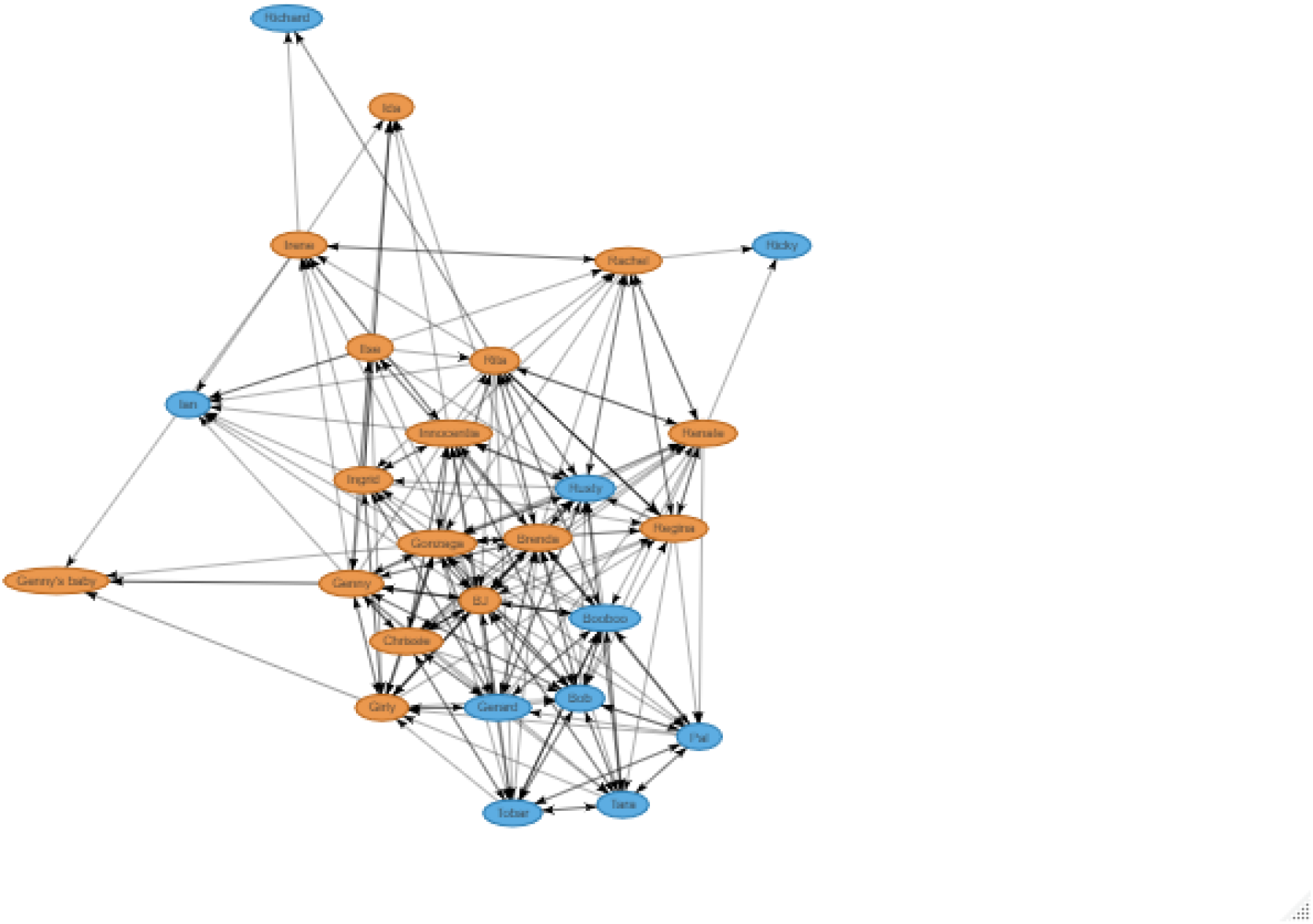
Grooming network - Group 1. Orange ovals represent females, blue ones represent males.

**Figure 5:**
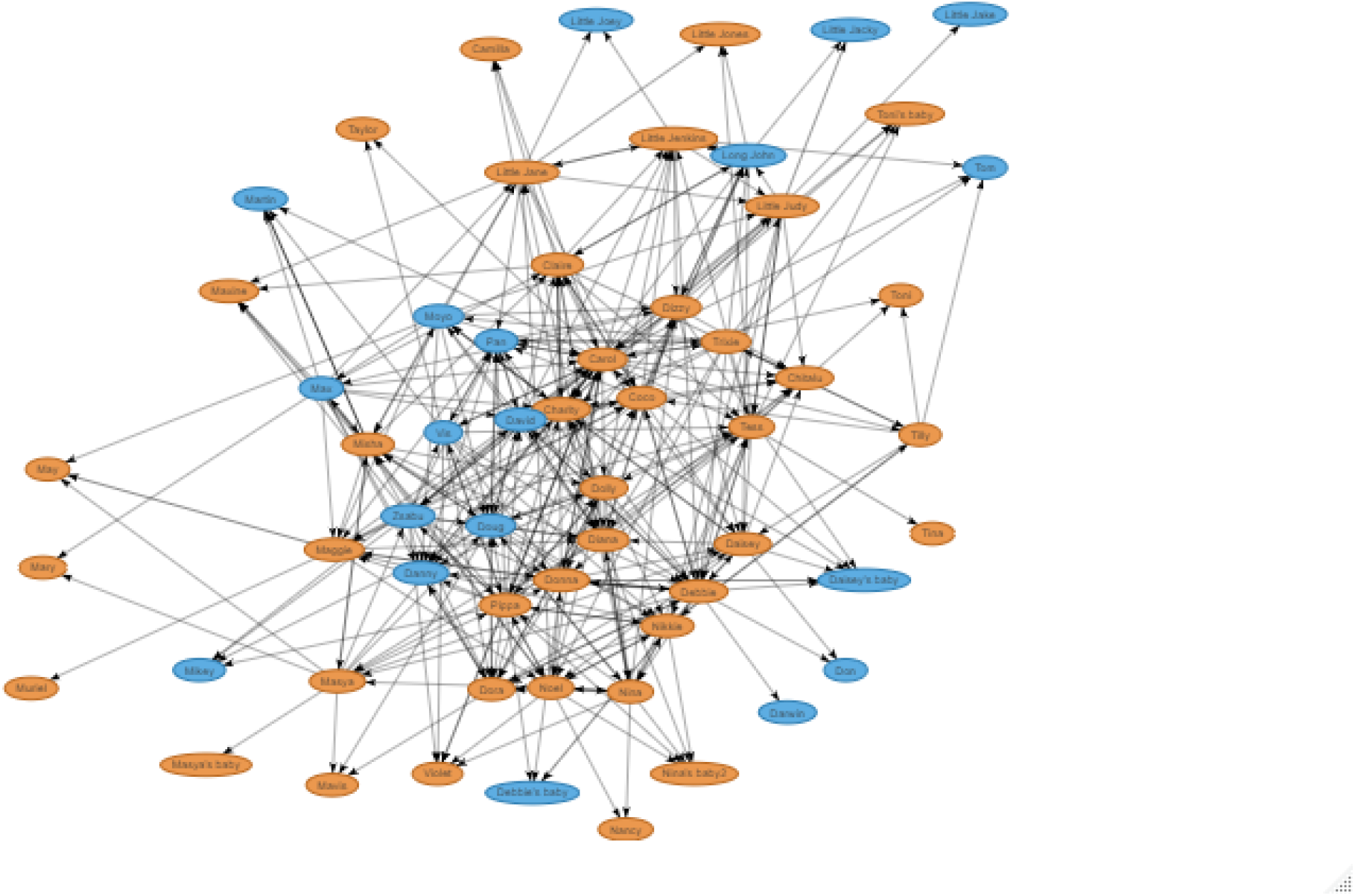
Grooming network - Group 2. Orange ovals represent females, blue ones represent males.

**Figure 6:**
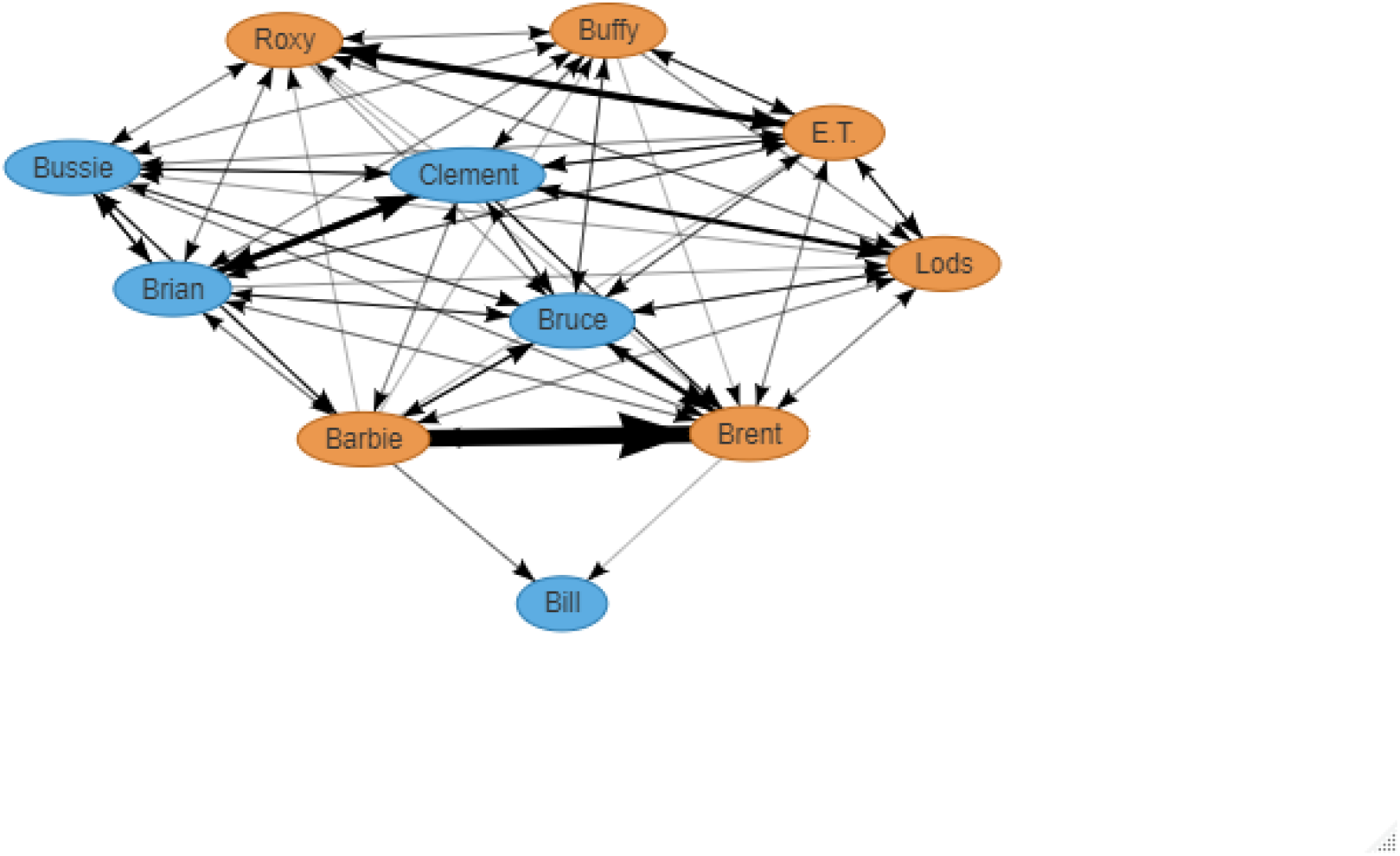
Grooming network - Group 3. Orange ovals represent females, blue ones represent males.

**Figure 7:**
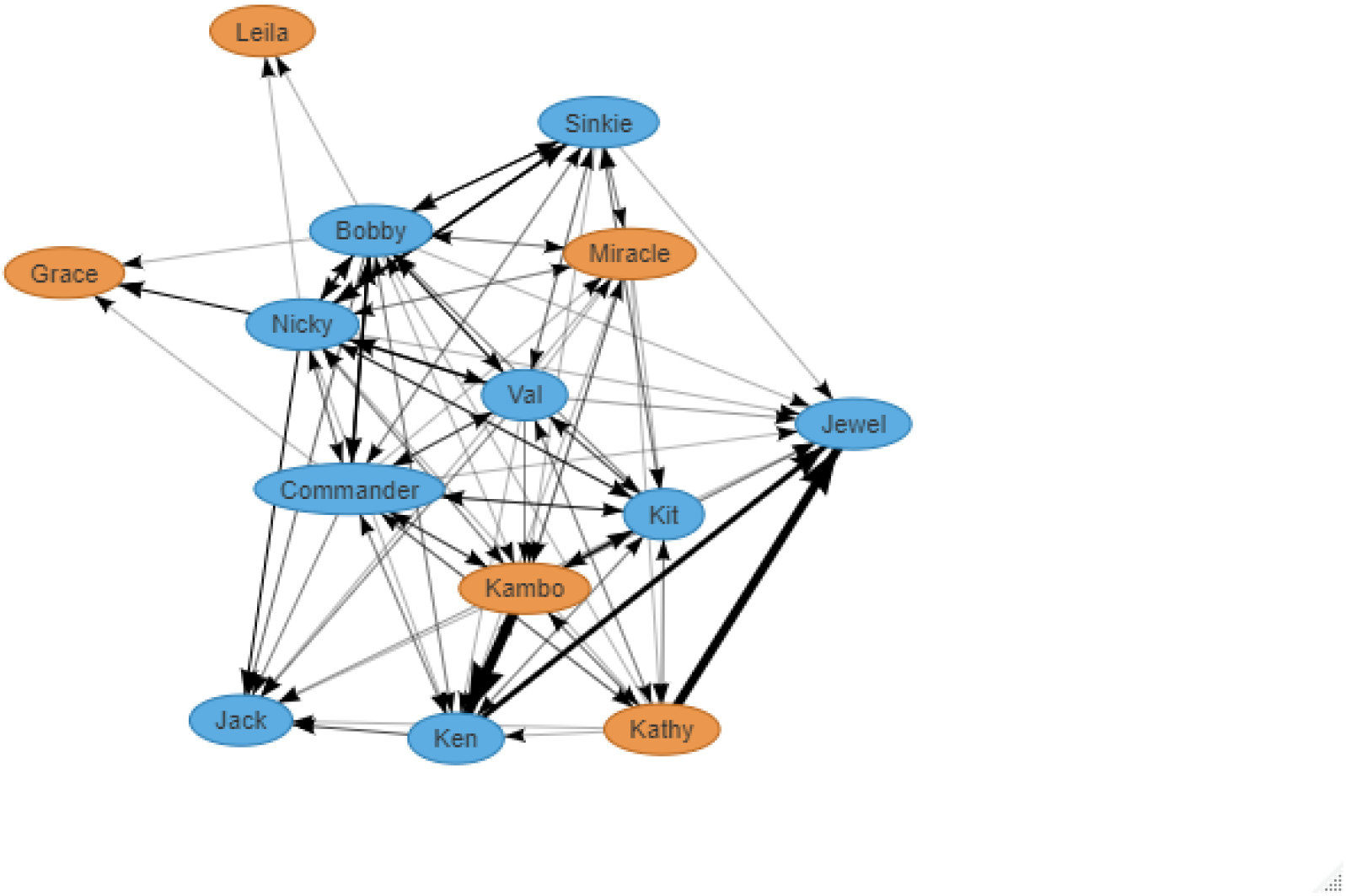
Grooming network - Group 4. Orange ovals represent females, blue ones represent males.

### Egonetworks

**Figure 8:**
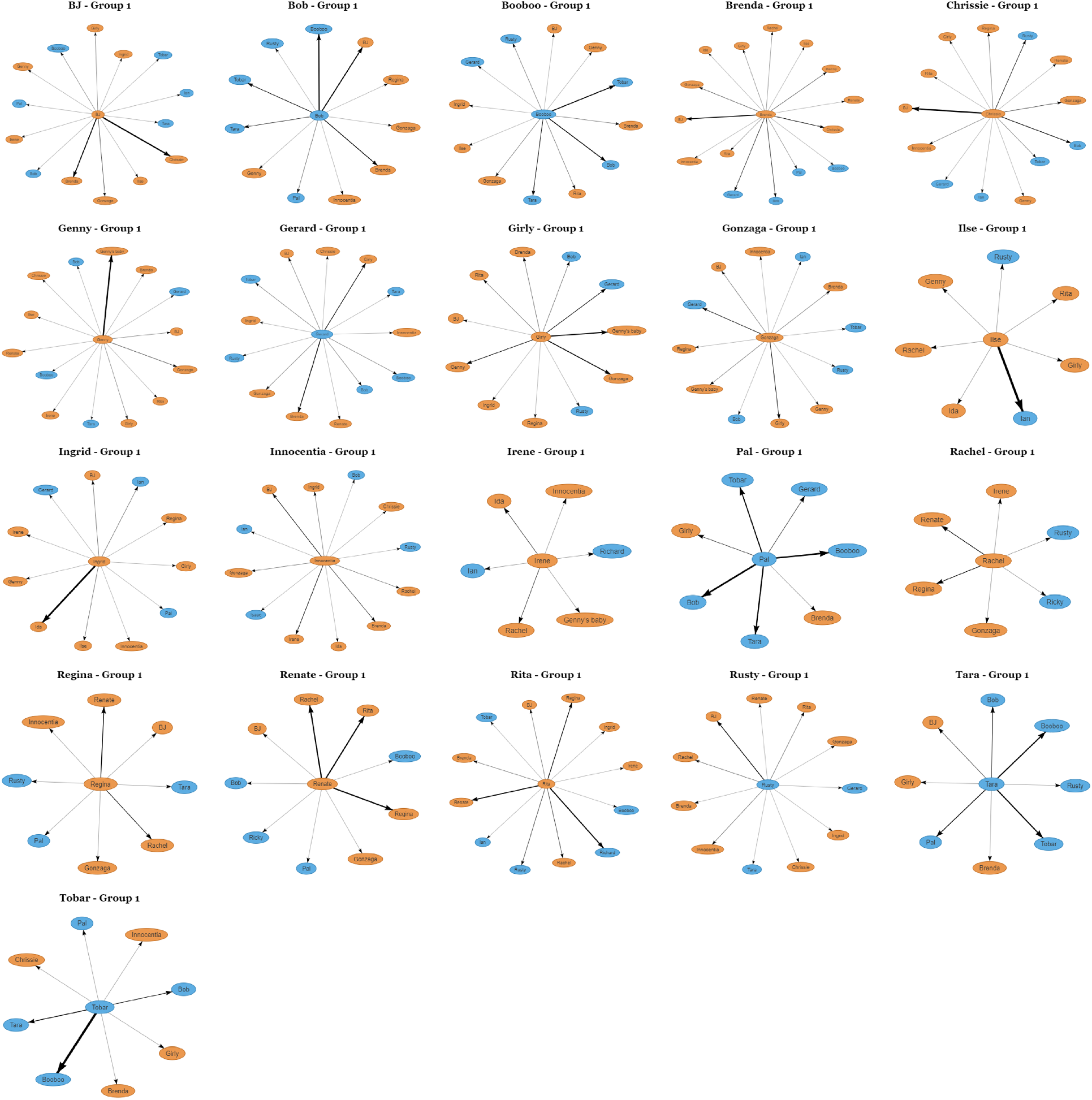
Egonetworks - Group 1. Orange ovals represent females, blue ones represent males.

**Figure 9:**
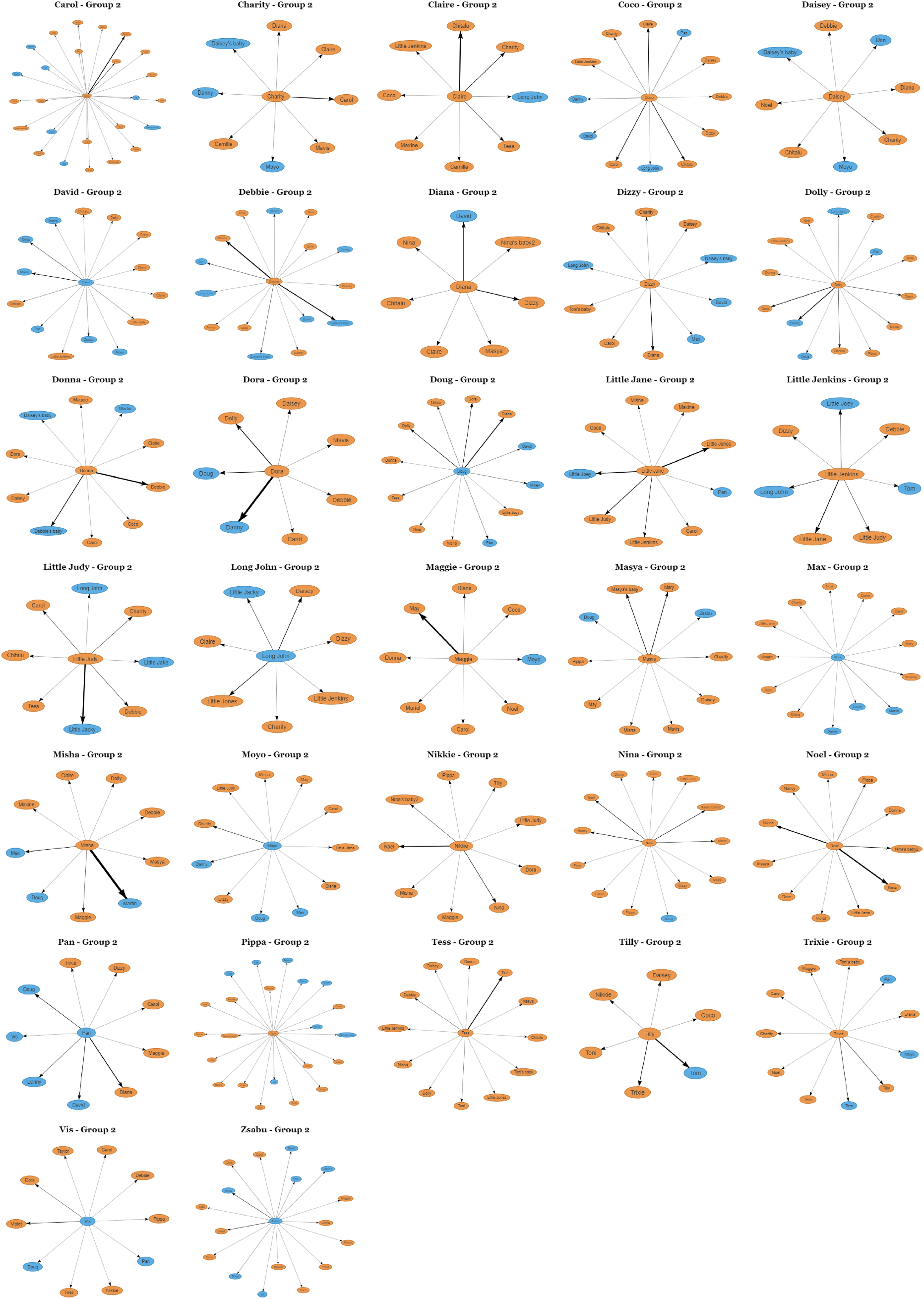
Egonetworks - Group 2. Orange ovals represent females, blue ones represent males.

**Figure 10:**
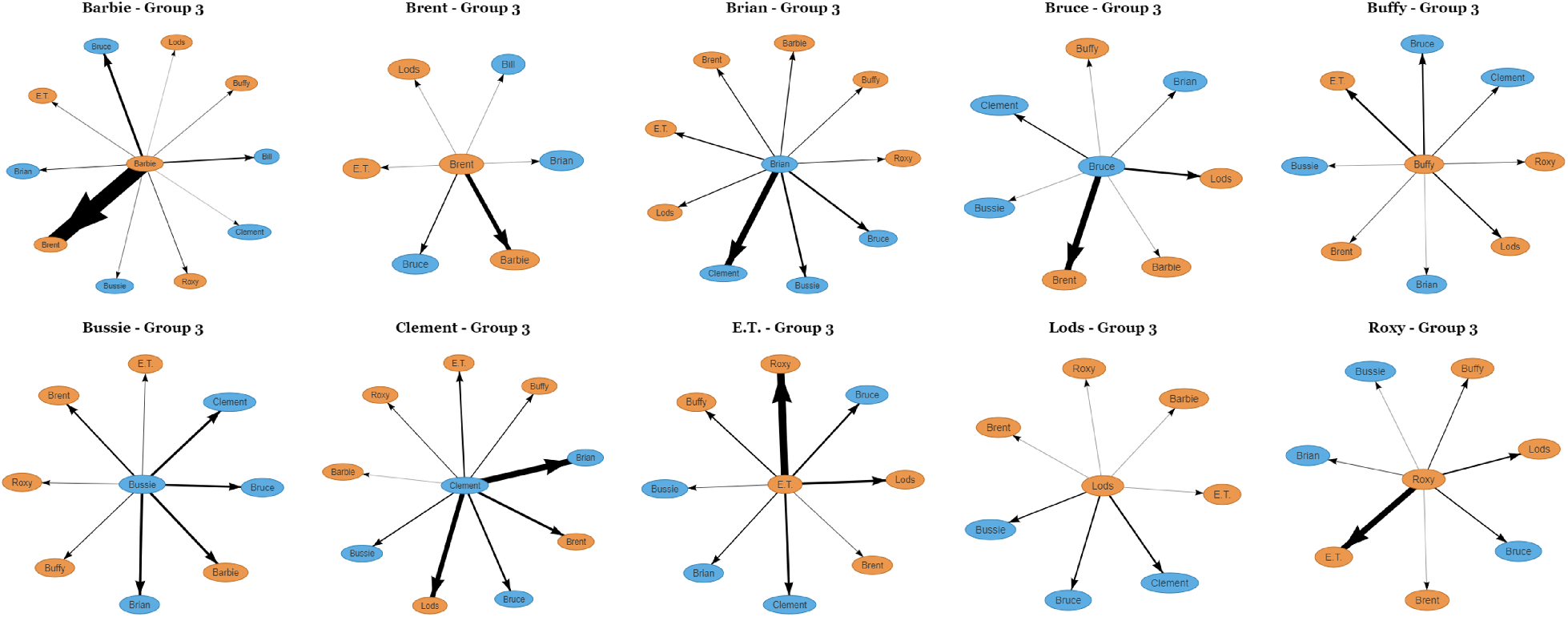
Egonetworks - Group 3. Orange ovals represent females, blue ones represent males.

**Figure 11:**
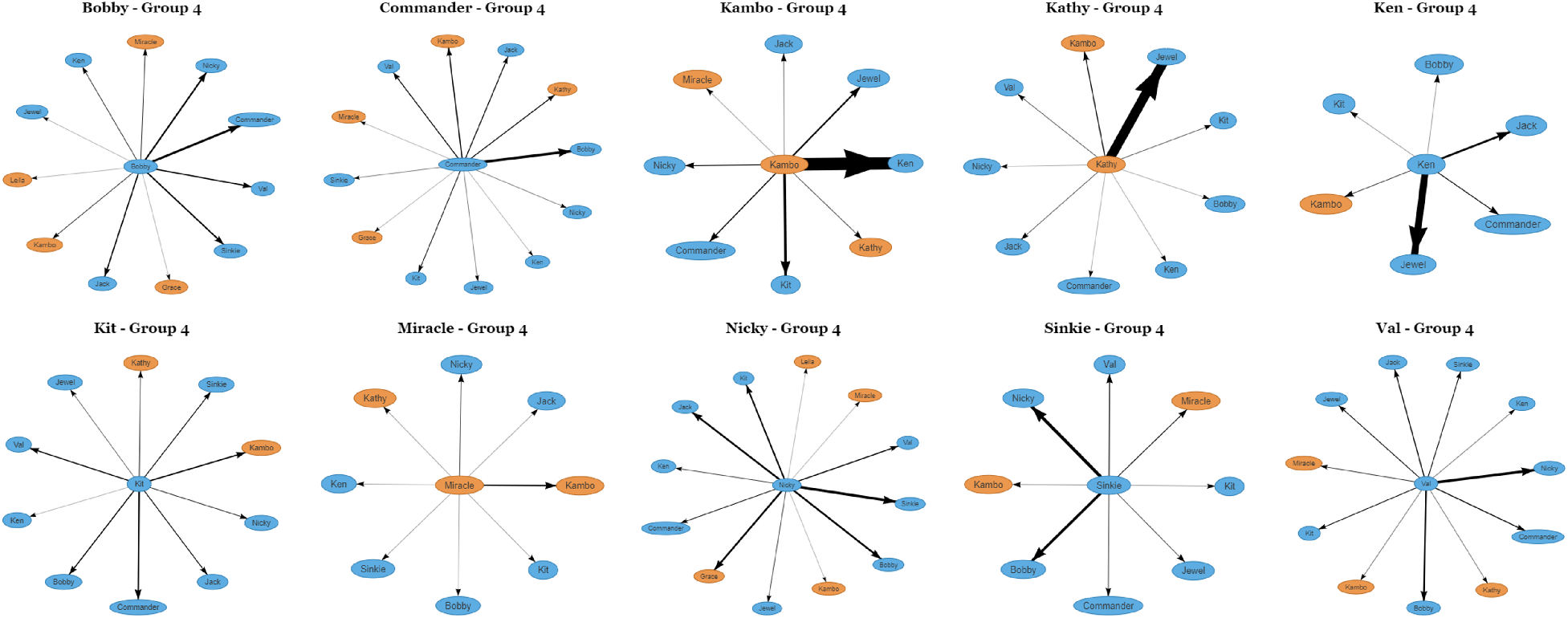
Egonetworks - Group 4. Orange ovals represent females, blue ones represent males.

### Individual fits of the distribution of ties

**Figure 12:**
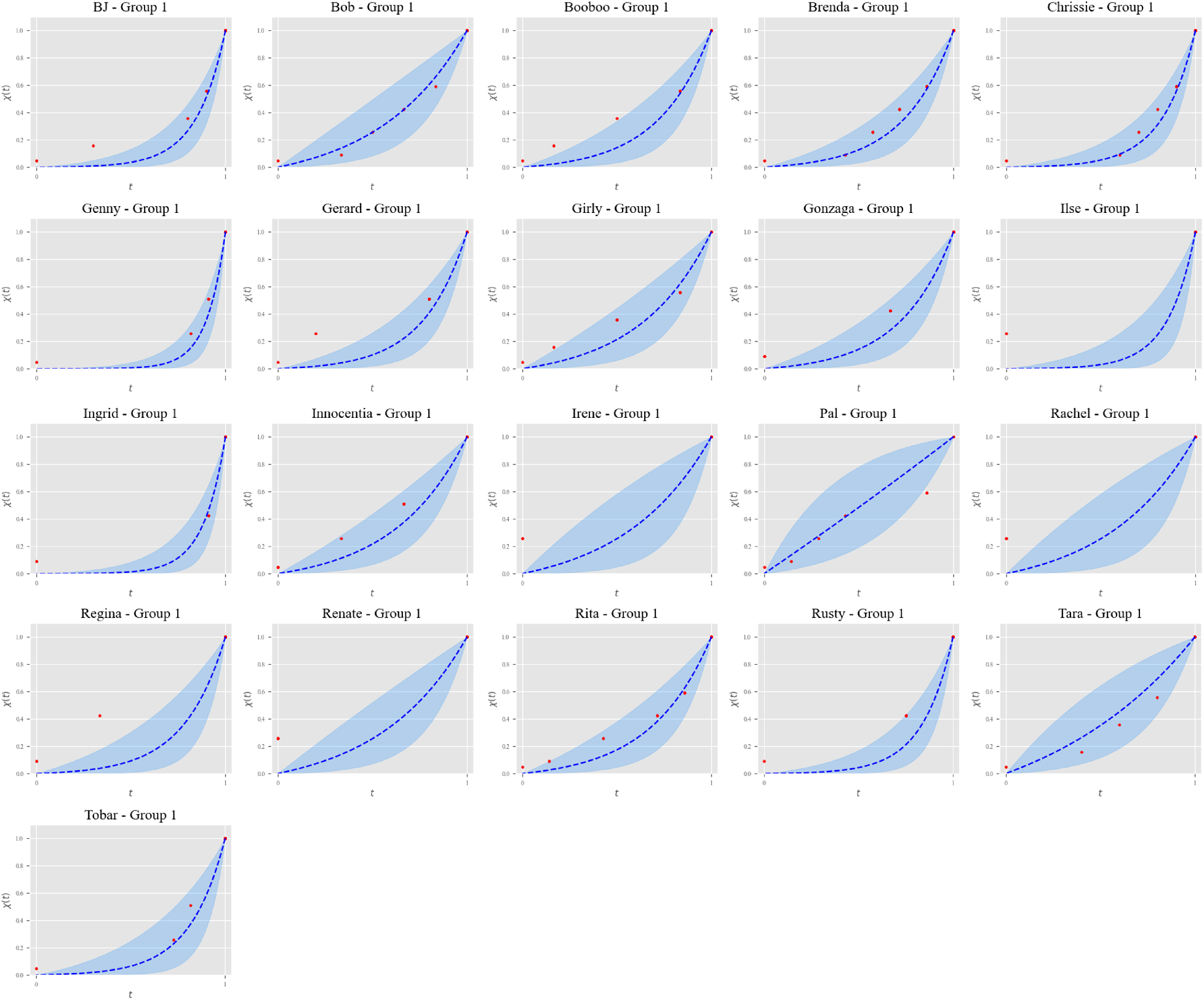
Individual fits of the distribution of ties - Group 1

**Figure 13:**
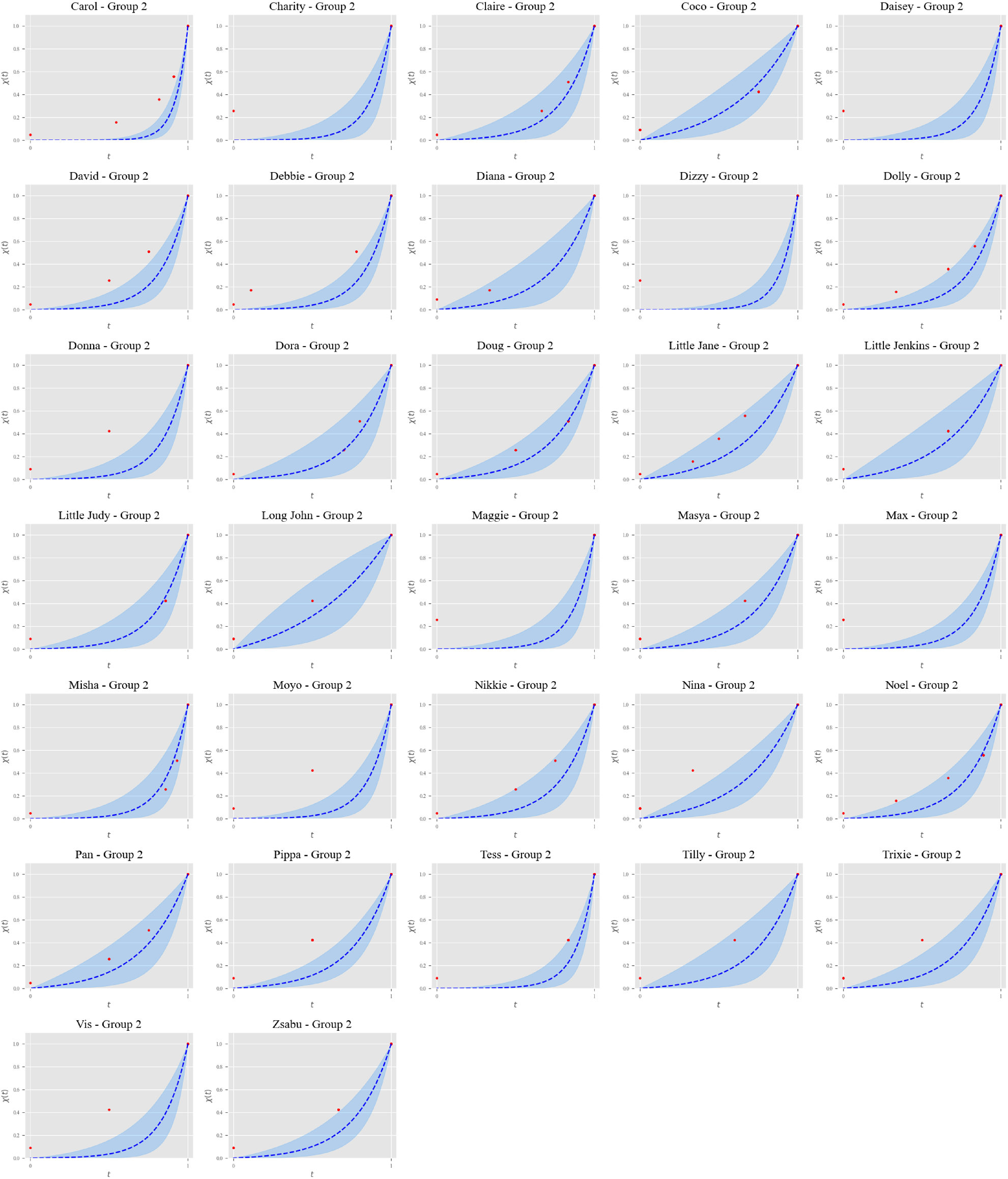
Individual fits of the distribution of ties - Group 2

**Figure 14:**
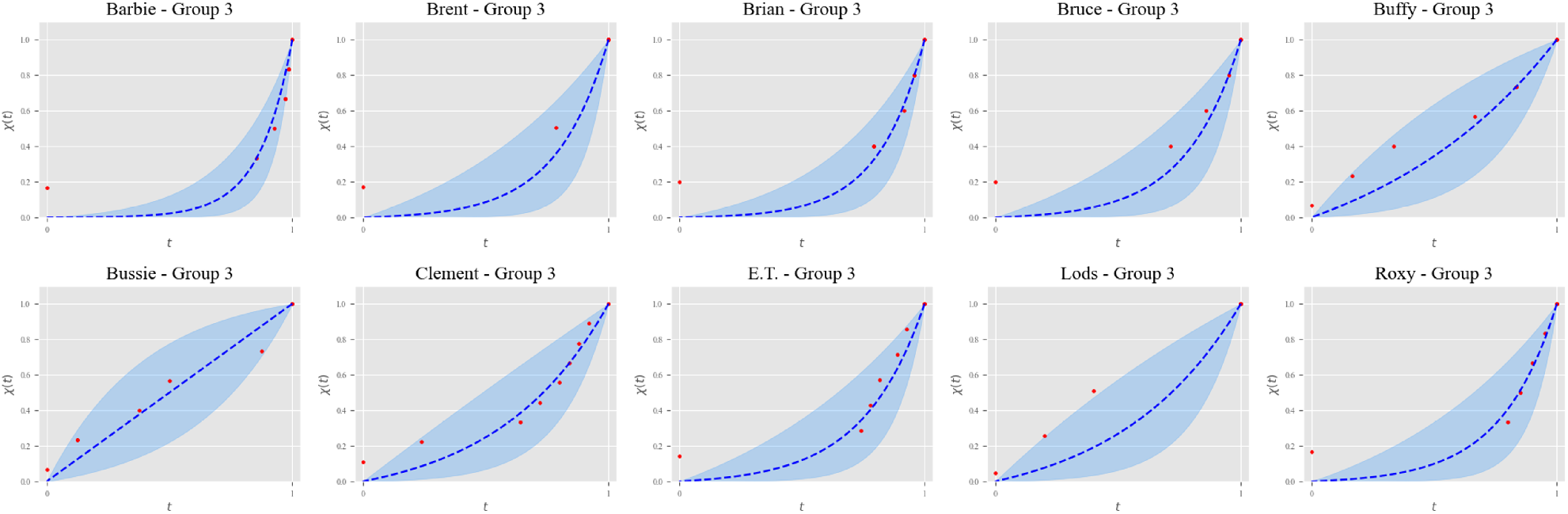
Individual fits of the distribution of ties - Group 3

**Figure 15:**
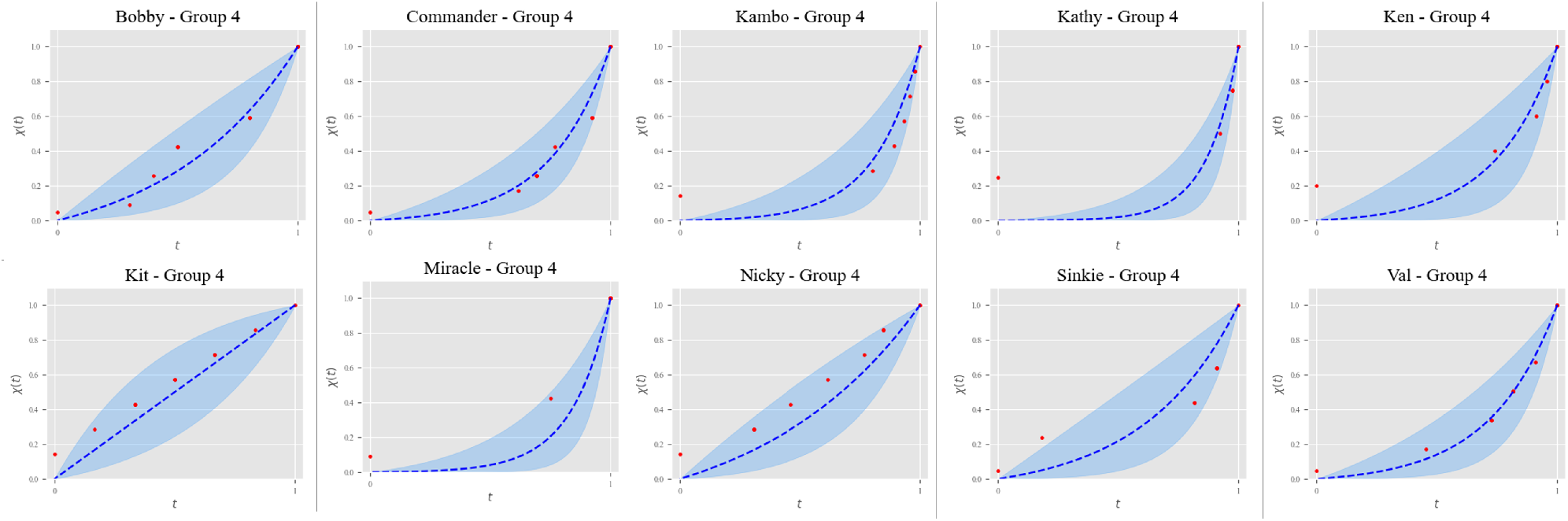
Individual fits of the distribution of ties - Group 4

## Acknowledgments

We are thankful to Robin Dunbar and José L. Molina for discussions on our results. DE, VD-M, JAC and AS acknowledge support from project BASIC (PGC2018-098186-B-I00) funded by MCIN/AEI/ 10.13039/501100011033 and by ‘ERDF A way of making Europe’. EJCvL acknowledges support from the Flanders Research Foundation (FWO) in the capacity of a postdoctoral fellowship. DBMH is supported by the Max Planck Society for the Advancement of Science.

## Author contributions

DE, VD-M, JAC and AS designed research, KAC, DBMH and EJCvL contributed the Chimfunshi data, DE and KAC curated the data, DE and VD-M analyzed the data, and all authors discussed and interpreted the results and wrote the manuscript.

## Data availability

Data are available from KAC, DBMH and EJCvL upon reasonable request.

## Materials & Correspondence

Correspondence and material requests should be addressed to AS.

